# Within-host infectious disease models accommodating cellular coinfection, with an application to influenza

**DOI:** 10.1101/359067

**Authors:** Katia Koelle, Alex Farrell, Christopher Brooke, Ruian Ke

## Abstract

Within-host models are useful tools for understanding the processes regulating viral load dynamics. While existing models have considered a wide range of within-host processes, at their core these models have shown remarkable structural similarity. Specifically, the structure of these models generally consider target cells to be either uninfected or infected, with the possibility of accommodating further resolution (for example, cells that are refractory to infection and cells that are in an eclipse phase). Recent findings, however, indicate that cellular coinfection is the norm rather than the exception for many viral infectious diseases, and that cells with high multiplicity of infection are present over at least some duration of an infection. The reality of these cellular coinfection dynamics is not accommodated in current within-host models although it may be critical for understanding within-host dynamics. This is particularly the case if multiplicity of infection impacts infected cell phenotypes such as their death rate and their viral production rates. Here, we present a new class of within-host disease models that allow for cellular coinfection in a scalable manner by retaining the low-dimensionality that is a desirable feature of many current within-host models. The models we propose adopt the general structure of epidemiological ‘macroparasite’ models that allow hosts to be variably infected by parasites such as nematodes and host phenotypes to flexibly depend on parasite burden. Specifically, our within-host models consider target cells as ‘hosts’ and viral particles as ‘macroparasites’, and allow viral output and infected cell lifespans, among other phenotypes, to depend on a cell’s multiplicity of infection. We show with an application to influenza that these models can be statistically fit to viral load and other within-host data, that they can reproduce notable features of within-host viral dynamics, and that important *in vivo* quantities such as the mean multiplicity of cellular infection can be easily quantified with these models once parameterized. The within-host model structure we develop here provides an alternative approach for modeling within-host viral load dynamics and allows for a new class of questions to be addressed that consider the effects of cellular coinfection, collective viral interactions, and viral complementation in within-host viral dynamics and evolution.

## Introduction

In part through the development and analysis of mathematical models, the processes driving the within-host dynamics of viral infections have been increasingly well understood over the last two decades. Statistical fitting of models to within-host data such as viral load measurements and immune response data have yielded estimates of within-host basic reproduction numbers for various viral pathogens, including HIV [1], influenza [2–4], measles [5], and dengue [6,7]. These fits have further characterized the roles of the innate immune response [2,3,7], and, particularly in secondary infections, the adaptive immune response [7,8] in regulating within-host viral dynamics. The structure of these within-host models have generally mirrored the structure of epidemiological ‘microparasite’ models, with cells being considered either uninfected or infected. In some models [2,3], uninfected cells have been further categorized as either susceptible or refractory to infection, again, mirroring hosts who are either susceptible or immune to infection in epidemiological models.

While these within-host models capture many of the important features of within-host viral processes, the majority of them implicitly assume that cellular coinfection does not occur [9] or that cellular coinfection, if it occurs, does not affect the phenotypes of infected cells [10,11]. Yet several experimental findings point towards cellular multiplicity of infection having the potential to impact cellular phenotypes such as the rate at which infected cells produce viral output [12,13] and the probability of initiating an interferon response [14]. The implicit assumption that a cell’s multiplicity of infection does not impact its phenotypes is hard-wired into ‘microparasite’-structured models because these models generally only consider a single class of infected cells, regardless of cellular multiplicity of infection. With increasing genomic evidence that cellular coinfection frequently occurs in chronic viral infections such as HIV [15] and HCV [16], as well as in acute viral infections such as influenza [17–19], a few notable models have been developed that have accommodated the possibility of cellular coinfection [10,11,20–22]. However, these models either remain high dimensional [11] or have made the assumption that host cell resources are limiting, such that viral output is independent of the extent of cellular coinfection [20]. While this assumption may be warranted for some viruses, it is likely not met in the case of many other viral pathogens.

Here, we develop a new class of low-dimensional within-host models whose structure flexibly allows for cellular coinfection. We base this new class of models on the structure of epidemiological ‘macroparasite’ models [23]. Development of these powerful epidemiological models started in the 1970’s [24,25], and they are now being commonly used to study how macroparasites such as nematodes spread through host populations [26]. They have further been used to assess the effect of control strategies on disease burden and host mortality [27]. We specifically develop this class of within-host ‘macroparasite’ models in the context of acute viral infections, although their structure can also easily accommodate the within-host dynamics of chronic infections. Finally, to demonstrate the usability of these models, we fit specific instances of these models to within-host equine influenza data.

## Methods

The structure of the within-host viral dynamic models we propose is based on a close analogy to population-level macroparasite models that are well-established and frequently used in the field of disease ecology and epidemiology (Figure 1). In Supplemental Section ‘Epidemiological macroparasite models’, we briefly review the derivation of the canonical structure of these population-level macroparasite models. Co-opting this canonical formulation for within-host viral dynamics allows us to flexibly model cells that have become infected with 0, 1, … *n* viral particles, while maintaining a low-dimensional set of equations.

**Figure 1.**
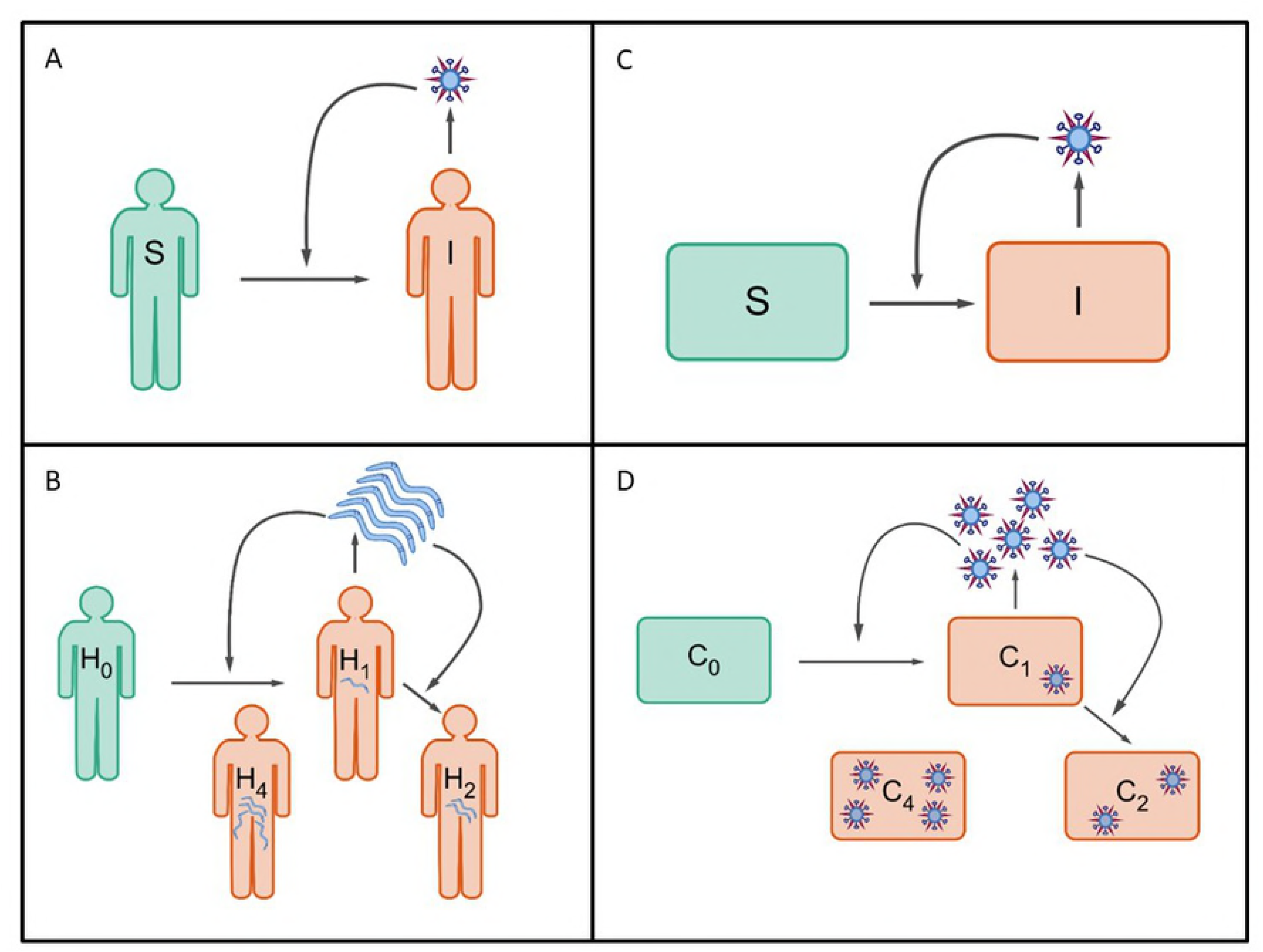
A schematic showing parallels between epidemiological and within-host infectious disease models. Epidemiological models fall into two groups: (A) models for microparasites and models for macroparasites, such as nematodes. Models for microparasites categorize individuals as being infected or uninfected. Models for macroparasites consider the parasite burden of infected individuals, as this burden affects the production rate of macroparasites from infected hosts and the mortality rate of hosts. (C) General structure of current within-host disease models. These models generally categorize cells as being infected or uninfected. (D) Schematic of a within-host ‘macroparasite’ model, proposed here. Models of this type would consider the multiplicity of cellular infection, as multiplicity of infection affects the rate of viral production and the lifespan of infected cells, among other phenotypes.

### Target-cell limited ‘macroparasite’ models

The simplest version of the within-host ‘macroparasite’ model is a target-cell limited model. In its most general form, this model is given by:

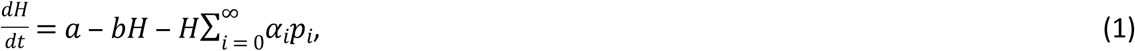

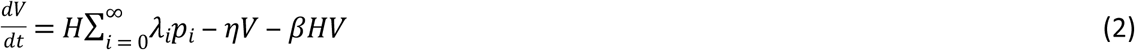

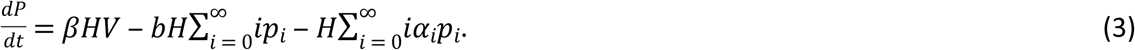

The variable *H* quantifies the total number of target cells, which includes both uninfected and variably infected cells. In this model, both uninfected and infected cells can be targets of further infection, so this variable differs from the variable representing uninfected target cells in traditional within-host ‘microparasite’ models. In equation (1), *a* is the constant rate of target cell production and *b* is the per capita background mortality of target cells. In the absence of infection, the target cell population equilibrates to *H*=*a*/*b*. In an acute viral infection such as influenza, the rate of target cell production *a* and the background rate of cell mortality *b* are frequently assumed to be sufficiently small to be ignored. The third term, 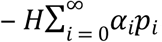, is the decrease in the number of target cells due to virus-induced mortality, with *p_i_* being the proportion of target cells that are infected with a cellular multiplicity of infection (MOI) of *i*.

The variable *V* quantifies the amount of free (extracellular) virus, and is analogous to the free virus variable generally modeled in traditional within-host ‘microparasite’ models. The first term, 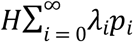, quantifies the overall rate at which free virus is produced from target cells, where *λ_i_* is the rate at which cells infected with an MOI of *i* produce free virus. The second term quantifies the rate of viral clearance, and the third term captures loss of free virus from its entry into target cells. This third term is often lost in traditional within-host ‘microparasite’ models, with an argument that loss of free virus from cell entry is negligible relative to loss of free virus through viral clearance [9].

The variable *P* quantifies the total amount of internalized virus across all target cells *H* and does not have an analogue in traditional within-host ‘microparasite’ models. This variable is related to, but distinct from, cellular multiplicity of infection (MOI), or equivalently, ‘cellular input’. While cellular MOI quantifies the number of virions a single cell has internalized, the variable P quantifies the number of internalized virions across all existing target cells. Mathematically, *P* can also be written as the product of the mean cellular MOI and the number of target cells *H*.

The first term in equation (3) captures the increase in the number of internalized virions from the entry of free virus *V* into target cells *H*. The second term, 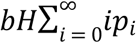, captures loss of internalized virus through background mortality of target cells, with the loss of a cell with an MOI of *i* leading to the loss of *i* internalized virions. The third term, 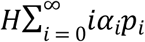, captures the loss of internalized virus through virus-induced mortality, with, again, the loss of a cell with an MOI of *i* leading to the loss of *i* internalized virions.

At this point, we can further simplify the model by adopting assumptions that may or may not be empirically supported. For instance, we can make analogous assumptions to the ones made in epidemiological ‘macroparasite’ models, e.g., that the viral production rate is linearly related to cellular MOI (*λ_i_* = *λi*), and that the cell mortality rate scales linearly with cellular MOI (*α_i_* = *αi*). Applying these assumptions, equations (1–3) become the following for an acute infectious disease:

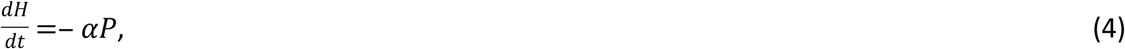

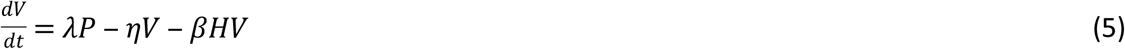

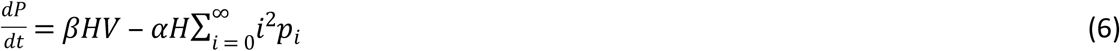

To further simplify equation (6), we can adopt an analogous assumption to one that is present in epidemiological ‘macroparasite’ models (see Supplemental Section ‘Epidemiological macroparasite models’): that the distribution of cellular MOIs is given by a negative binomial distribution with mean *P*/*H* and dispersion parameter *k* in the range of (0, ∞). This assumption simplifies equation (6) to:

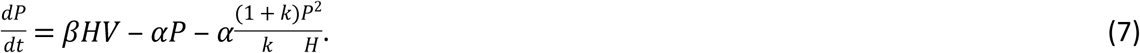

The negative binomial distribution allows for the possibility of cellular MOI overdispersion (low *k*), while still allowing for a Poisson distribution of cellular MOIs when *k* = ∞. Overdispersion of cellular MOIs *in vivo* is highly likely for several reasons. First, some target cells might be more susceptible to infection that others due to variation in the number and types of receptors. Second, given spatial aspects of within-host viral spread, there is likely considerable variation in the rate at which cells are exposed to virus. Third, variation in the time cells remain in their eclipse phase can under certain conditions produce overdispersion of cellular MOIs (see Supplemental Section ‘Negative binomial distribution for multiply infected cells’).

We can define the basic reproduction number *R*_0_ for the target-cell limited model given by equations (4), (5), and (7) as the number of new viral particles generated by a single viral particle at the onset of an individual’s infection when the overwhelming majority of target cells are uninfected. To derive *R*_0_ for this model, we can first make a fast viral dynamics assumption, such that 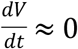 and 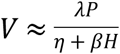. Plugging this expression into equation (7), and recognizing that the ratio 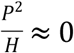 at the onset of infection, yields: 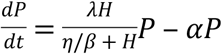. From this expression, it is clear that 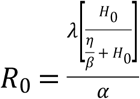, where *H*_0_ is the number of target cells present at the beginning of the infection.

While we assume fast viral dynamics in the derivation of *R*_0_, we model within-host viral dynamics under the target-cell limited version of this model using all three variables (*H, V*, and *P*) because it is uncommon to assume fast viral dynamics in within-host models and retaining *V* in the model allows for a more straightforward interface with viral load data.

Since little is known about how viral production rates and cellular mortality rates scale with cellular input, alternative assumptions can also be made that would still allow for a simplification of equations (1)-(3). For example, it could be assumed that viral production rates are independent of cellular input, as long as a cell is infected. This assumption would implicitly assume that host cell machinery is the limiting factor governing viral production from a cell. This assumption would lead to equation (2) becoming:

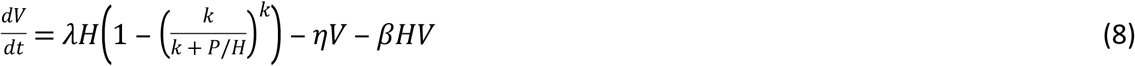

where the term 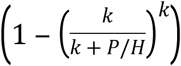 provides the probability that a target cell has internalized at least one viral particle. It could also be assumed that the cellular mortality rate is independent of cellular input (as long as there is some input), and equations (1) and (3) would be modified accordingly:

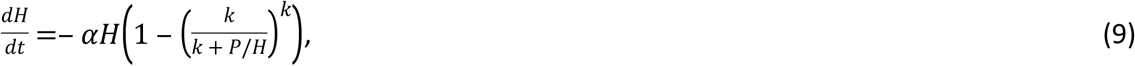

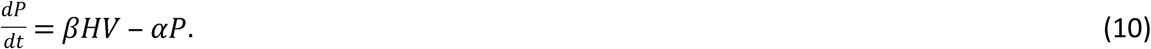

Under the assumption that virions are internalized independently of a cell’s MOI, the dispersion parameter *k* = ∞ and equations (8)-(9) become:

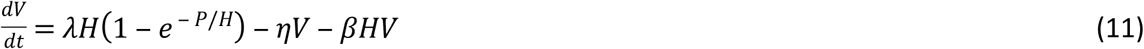

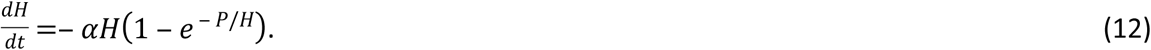

Defining the number of currently infected target cells as *I* = *H*(1 ‒ *e* ‒ *P*/*H*) allows one to expand equation (12) into uninfected (*T*) and infected (*I*) target cell classes: 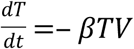 and 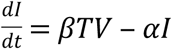, respectively. Neglecting the third term, this definition also allows us to simplify equation (11) to 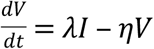. The variable *P* can be excluded if it assumed to be in equilibrium with *V*. As such, it is clear that with the assumption of Poisson-distributed cellular MOIs, an MOI-independent viral production rate, and an MOI-independent mortality rate of infected cells, the within-host ‘macroparasite’ model folds into the traditional within-host ‘microparasite’ model. This finding is consistent with findings from a previous, high dimensional model for HIV that accommodated multiply-infected cells. An analysis of that model showed that it simplified to the structure of a within-host ‘microparasite’ model that had all infected cells belonging to a single infected class *I* when viral production rates (and infected cell mortality rates) were independent of the number of internalized virions [20].

The ‘true’ relationship between cellular input and the rate of viral production likely depends on virus and host cell type, and needs to be empirically addressed when applying the within-host ‘macroparasite’ model to a specific viral infection. Similarly, little is known about how cellular mortality rate scales with cellular input and experimental studies need to be performed to address this knowledge gap.

### Within-host ‘macroparasite’ models incorporating the host’s immune response

Within-host models of viral infections frequently incorporate the host’s immune response, since it is this response that is thought to play a critical role in regulating and ultimately clearing viral infections [7,25]. Minimally, the host’s immune response can be incorporated by considering only the innate immune response, which can, again at minimum, be captured by a single additional variable. This variable can encompass the activity of interferons and cytokines, as well as cells of the innate immune response such as natural killer (NK) cells. In many models, the dynamics of specifically interferon-*α* have been included, with interferon production occurring from infected cells and decaying at a constant rate [2,3,7]. If we assume that cells produce interferon at a rate (or probability) that scales linearly with cellular MOI, then the dynamics of interferon-*α* are given by:

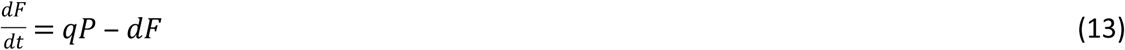

Interferon-*α* can modify viral within-host dynamics in a number of ways. One way is for interferon-*α* to reduce the rate of viral production from infected cells. Another way is for interferon-*α* to decrease the susceptibility of cells to infection (or further infection). Both of these mechanisms of action can be assumed to respond to immediate levels of interferon. In this case, the viral production rate can be reduced from 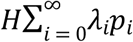 (equation 2) to 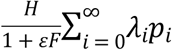, or similar (as in [28]), and the rate of viral entry into target cells can be reduced from *βHV* (equations 2 and 3) to 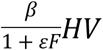 or similar (as in [3]). Alternatively, cellular exposure to interferon could have prolonged effects, with cells becoming refractory to infection (or further infection) for a period of time (as in [2,3]) and infected cells reducing their viral output for a period of time following exposure. A third, indirect effect of interferon-*α* is to facilitate the recruitment of innate effector cells, which would act to clear infected target cells, leading to an overall effective increase in the rate at which target cells decline. Here, for simplicity, we consider the two direct effects of interferon: the effect of these molecules on reducing cell susceptibility to infection and on reducing the rate of viral production from infected cells, assuming that interferon has prolonged effects on cells. Our innate immune response model is given by equation (13) and the following system of equations:

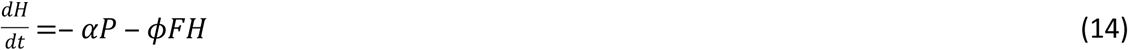

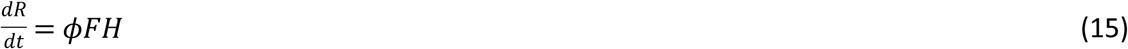

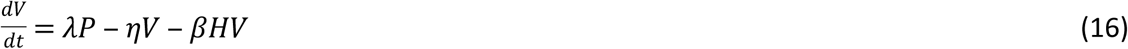

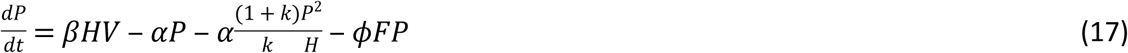

Here, *H* is the total number of *susceptible* target cells (including infected and uninfected target cells), *R* is the total number of target cells that are *refractory* to further infection (including uninfected cells and already infected cells), *V* is again the amount of free virus, and *P* is the total number of viral particles across *susceptible* target cells. The parameter *ϕ* quantifies the per capita rate at which interferon makes cells refractory to infection. This model formulation assumes that all susceptible cells, whether uninfected or infected, become refractory to infection (or further infection) at similar rates, that refractory cells stay permanently refractory, and that no virus is produced from refractory cells. This latter assumption effectively reduces the overall rate at which the total infected cell pool produces virus as a result of interferon exposure. Model equations (13–17) assume analogous effects of interferon as the within-host ‘microparasite’ model presented in [3], while adopting assumptions of linear scaling between cellular MOI and infected cell mortality rate, between cellular MOI and viral production rate, and between cellular MOI and the probability of cellular interferon production.

## Results

Here, we fit the within-host macroparasite models presented above to empirical within-host data, highlighting biological insight that can be gained from these models that within-host ‘microparasite’ models have difficulty providing. We first fit the target cell limited model given by equations (4), (5), and (7) to influenza A/H3N8 viral load measurements from experimentally infected ponies. Prior to fitting this model, we confirmed that all of its parameters are structurally identifiable (Supplemental Section ‘Identifiability analysis’). However, due to the limited number of data points for each pony, we did not attempt to estimate all model parameters for each pony independently. Instead, based on previous analyses of these data [2,3], we set the initial number of target cells (in our case, given by the variable *H*) to 3.5 x 10^11^ cells, and further set the initial number of internalized virions *P* to 0. This latter assumption corresponds to the assumption made in previous models of 0 initially infected cells. We further constrained the per-particle production rate *λ*, the per-particle cellular mortality rate *α*, and the dispersion parameter *k* to be the same across infected ponies. We let the viral clearance rate *η*, the viral infectivity rate *β,* and the initial amount of free virus *V*(*0*), differ across ponies, since in part these parameters reflect host-specific characteristics or phenotypes related to a host’s immune history. Under these constraints, all parameters of this basic model were practically identifiable.

To estimate model parameters, we used a maximum likelihood approach that further accommodates below the limit of detection measurements (Supplemental Section ‘Statistical estimation of model parameters’). Figure 2a shows the viral load measurements from the ponies, along with the fit of the target cell limited macroparasite model (equations 4, 5, and 7). Table 1 lists the estimated model parameters. The model reproduces key features of the observed within-host influenza dynamics. Most notably, the within-host macroparasite model can hit peak viral loads and reproduce the biphasic viral declines that are observed in several of the ponies. This stands in contrast to the within-host, target-cell limited ‘microparasite’ model (for example, equations 1–3 in [4]), which cannot reproduce the biphasic viral decline and has trouble hitting peak viral loads.

**Figure 2.**
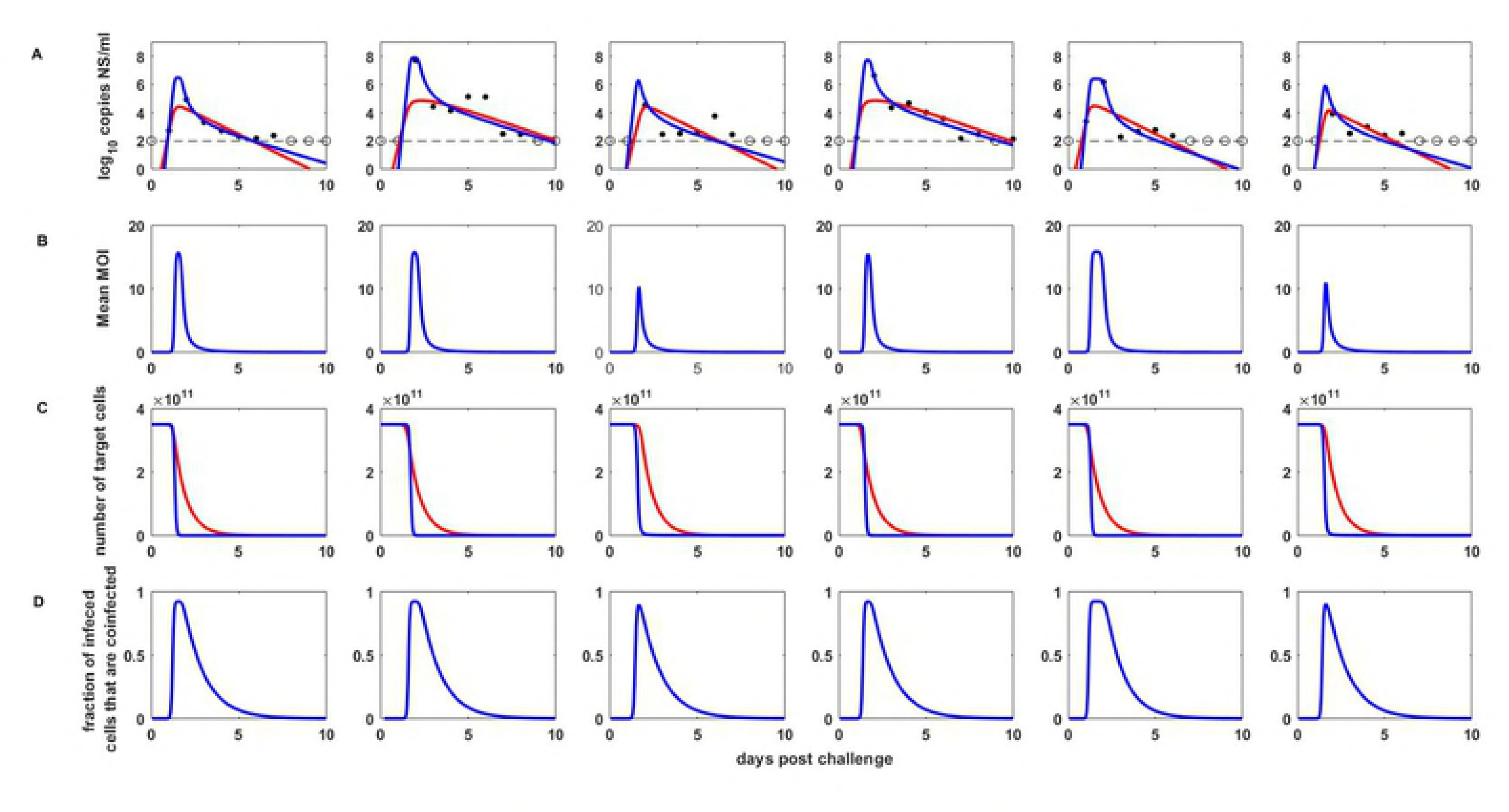
Target-cell limited within-host macroparasite model dynamics. (A) Within-host viral dynamics, parameterized by fitting the model to influenza A/H3N8 viral load measurements from experimentally infected ponies (black dots). The dashed black line shows the limit of detection, and open circles show below the limit of detection measurements. Blue lines show maximum likelihood fits of the within-host target-cell limited macroparasite model. Red lines show the maximum likelihood fits of a within-host target-cell limited microparasite model (equations 1–3 in [4]). The macroparasite model fits the data considerably better than the microparasite model (log likelihood = −49.76 vs. −57.87, with the same number of estimated parameters). (B) Mean multiplicity of infection over time for each of the ponies shown in (A), as calculated from the within-host ‘macroparasite’ model. Mean multiplicity of infection is calculated as the total number of intracellular particles divided by the total number of target cells, *P*(*t*)/*H*(*t*), where *t* is time since infection. (C) The number of target cells over time for the within-host ‘macroparasite’ model (blue lines; *H*) and for the within-host ‘microparasite’ model (red lines; *T* + *I*). (D) The proportion of infected cells that are infected by more than one viral particle, calculated from the within-host ‘macroparasite’ model.

**Table 1.**
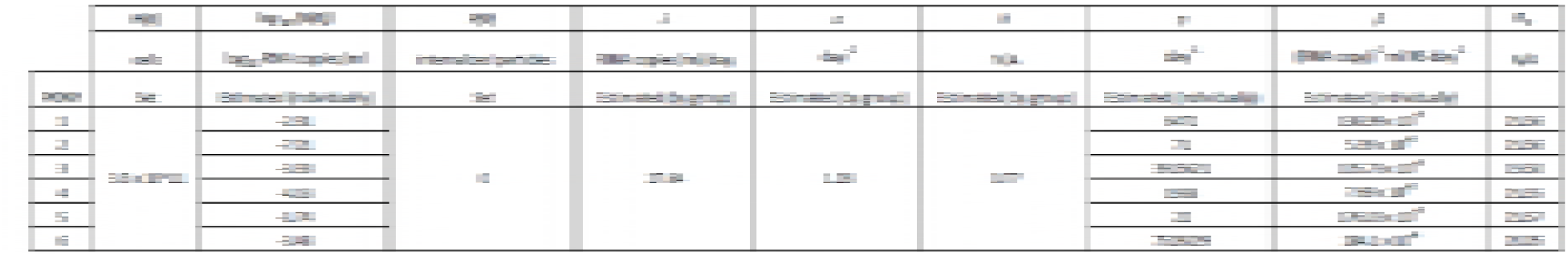
Parameter estimates for the target cell limited ‘macroparasite’ model, for each of the six ponies. The model is given by equations (4), (5), and (7). The within-host basic reproduction numbers for each pony are also listed, which are calculated from the listed parameter estimates according to the formula provided in the text.

In addition to being able to better reproduce these two key features of the viral dynamics than the within-host microparasite model, the within-host macroparasite model allows one to calculate the time-varying mean multiplicity of infection (Figure 2b). The mean multiplicity of infection in this model is simply given by the total number of internalized particles divided by the total number of target cells: *P*/*H*. Plotting the mean multiplicity of infection over time allows us to better understand the processes that are driving the biphasic viral declines projected by the macroparasite model. As viral load increases, the mean multiplicity of infection increases dramatically. Infected cells with high multiplicity of infection then experience high mortality rates, leading to a very rapid decline in viral load and a rapid depletion of target cells *H* (Figure 2c). Mean cellular MOI drops as a result of this rapid depletion of cells with high MOI (Figure 2b). It then remains low because of low levels of free virus *V* and thus little opportunity to internalize more virus. The second phase of the viral decline comes about from the low mortality rate of cells with low MOI.

One can also easily calculate the proportion of infected cells that are infected by more than one viral particle (Figure 2d). This information may be useful for anticipating the extent to which viral reassortment occurs *in vivo*, or for characterizing the landscape available to defective interfering particles, which ‘cheat’ off of wild-type virus for their own replication.

More recent quantitative analyses and data have indicated that influenza viral dynamics are likely regulated by the host immune response, particularly the innate immune response [2,3]. We therefore also fit the innate immune response model, provided by equations (13)-(17), to the available equine data, which now include viral load measurements, fold change measurements in interferon-*α,* and an estimate that only ~27% of host cells were depleted by the end of an infection (Supplemental Section ‘Statistical estimation of model parameters’; see [2,3] for descriptions of the data). Again, we constrained a subset of the model parameters to be the same across the ponies, while letting other parameters be pony-specific. Specifically, we constrained parameters *α, ϕ, λ, d*, and *k* to be the same across the ponies, because these parameters quantify infected cell phenotypes that we expect to be common across individuals. We let parameters *η, β, q*, and *V*(*0*) differ across ponies, due to host-specific factors. Figures 3a and 3b shows that this within-host macroparasite model can quantitatively reproduce features of the viral load and interferon-*α* measurements, without depletion of target cells to unreasonable levels (Figure 3c). Table 2 provides parameter estimates for this model. Here, the very rapid initial viral decline results from several processes: the cellular input-dependent mortality rate of infected cells, the rapid removal of susceptible target cells *H* through exposure to interferon-*α*, and the lack of viral production from refractory cells. The second, slower phase of viral decline results from the slow clearance of the remaining infected cells that have low multiplicity of infection.

**Figure 3.**
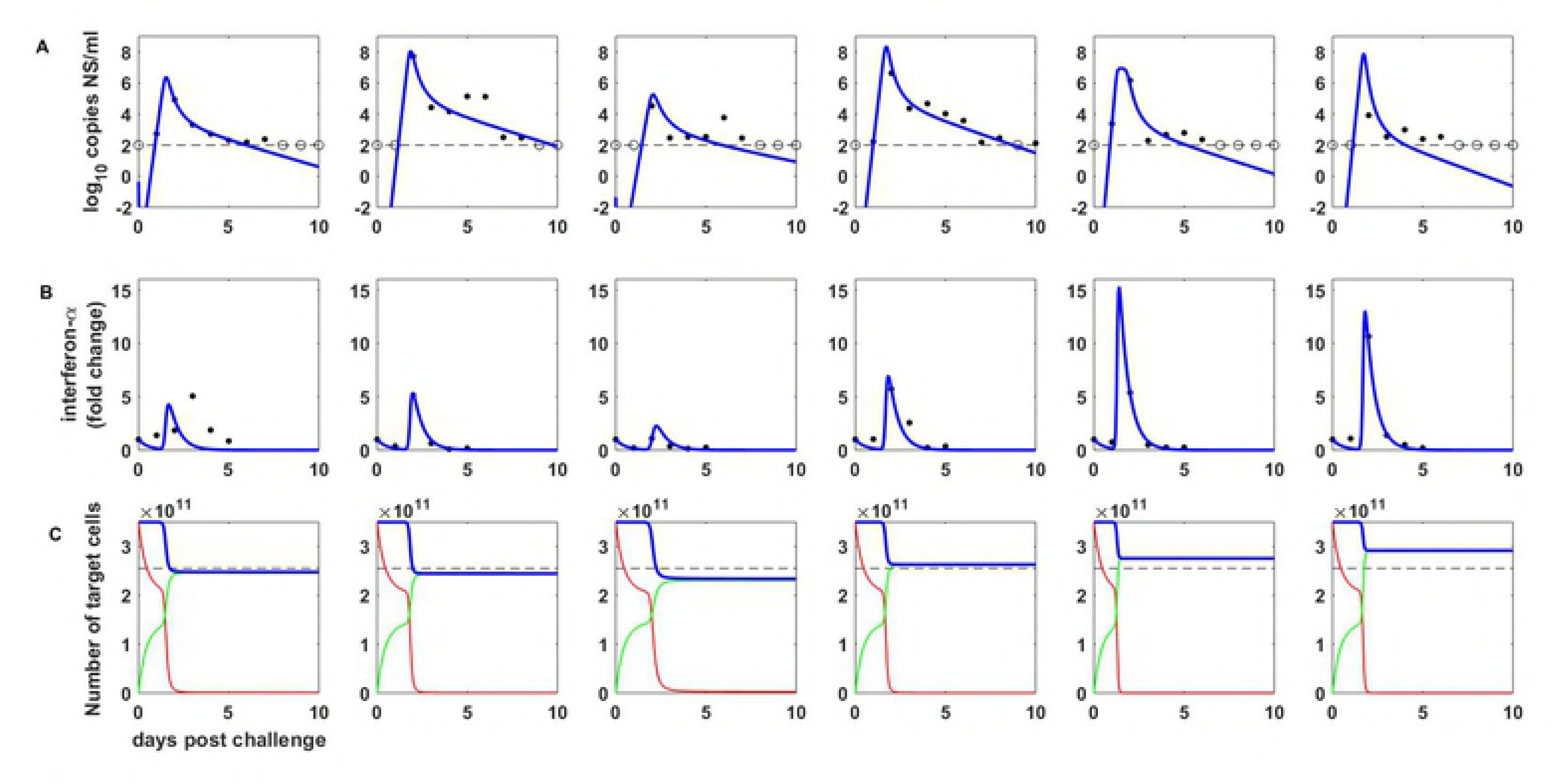
Within-host dynamics from the macroparasite model incorporating the innate immune response. The model, given by equations (13)-(17) was parameterized by fitting to influenza A/H3N8 viral load measurements (A) and fold change measurements of interferon-*α* (B) from experimentally infected ponies. We further used an estimate of 73% of target cells remaining at the end of infection (C) to parameterize the model. (A) Within-host viral dynamics, with data as in Figure 2A. Blue lines show maximum likelihood fits of the within-host macroparasite model incorporating the innate immune response. (B) Interferon-*α* dynamics. (C) Target cell dynamics. Red lines show the dynamics of susceptible target cells *H*, green lines show the dynamics of refractory target cells *R*, and dashed blue lines show the dynamics of all target cells (*H* + *R*). Dashed black line shows an estimate for the final number of target cells, given by a 27% reduction in the number of target cells [3].

**Table 2.**
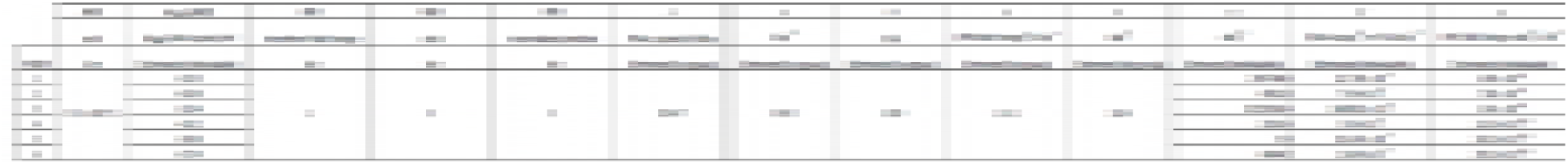
Parameter estimates for the ‘macroparasite’ model incorporating the innate immune response, for each of the six ponies. The model is given by equations (13)-(17).

## Discussion

Here, we have developed a new class of within-host models for understanding the *in vivo* dynamics of viral infections. This new class of models differs from existing within-host models in that it allows for target cells to be variably infected and for the degree of cellular input to impact the phenotypes of infected cells, such as their death rate and the rate at which they produce virus. Despite allowing for cellular coinfection dynamics, these models remain low-dimensional. This addresses existing concerns in the literature about the scalability of within-host models that allow for cellular coinfection [11]. While we have developed and applied these within-host models to acute viral infections, the structure of these models is immediately applicable to chronic viral infections. When applied to chronic infections, the terms we lost for the replenishment and natural death of target cells (see the Methods section) would simply need to be reintroduced.

The structure of the within-host models we derived here co-opt the structure of epidemiological ‘macroparasite’ models. Based on empirical data, those models generally assume that host death rates scale linearly with macroparasite burden and that the rate of egg release from infected hosts similarly scales linearly with macroparasite burden. The within-host models we fit similarly adopt these scaling relationship assumptions, although other scaling assumptions can be adopted while retaining the desirable low-dimensionality of the model equations. Clearly, the structure of the within-host model should reflect empirically supported relationships between cellular input and cellular phenotypes. To date, very few studies have attempted to empirically quantify these relationships, making the appropriate choice of model structure difficult to decide upon. For influenza, the studies that do exist have shown that the timing and amounts of viral yield depend critically on cellular input, and that cumulative viral yield generally increases with cellular input [12,13]. Intriguingly, these results stand in contrast to the current structure of the model that we have presented. This is because, if we assume that both the viral production rate and the cell death rate scale linearly with cellular input, the total cellular output of an infected cell should be independent of its multiplicity of infection *i*, with total cellular output being given by: *λi*/*αi* = *λ/α*. The empirically determined relationship between higher cellular output with higher cellular input therefore seems to indicate that cellular death rates must scale less than linearly with cellular input and/or that viral production rates must scale faster than linearly with cellular input. The latter relationship, if empirically supported, would provide tantalizing evidence for viral cooperation within cells playing a role in within-host viral dynamics, an idea that has recently gained traction [29–31].

A key feature of epidemiological macroparasite models is the possibility of nematode overdispersion across hosts. Parasite overdispersion has considerable empirical support, with the overwhelming majority of macroparasite distributions studied having a variance to mean ratio exceeding 1 and an estimated dispersion parameter *k* of less than 1 [32]. A more recent analysis further indicates that observed levels of parasite overdispersion can be attributed almost entirely to host heterogeneity in parasite exposure or host heterogeneity in susceptibility to infection [33]. Here, in the context of within-host viral dynamics, we also found statistical support for very high levels of overdispersion, with a dispersion factor *k* estimate of 0.77 in the target-cell limited model (Table 1) and an estimate of *k* = 0.30 in the innate immune response model (Table 2). This overdispersion could similarly reflect variation in target cell susceptibility to infection. It could also reflect heterogeneity in viral exposure, likely due to the intrinsically spatial aspect of influenza virus spread within infected hosts. Finally, as described in Supplemental Section ‘Negative binomial distribution for multiply infected cells’, overdispersion could also result from the distribution of time that cells remain in the eclipse phase prior to becoming productively infected. Regardless of the causes of viral overdispersion across target cells, the consequences of viral overdispersion are that the majority of infected cells are multiply infected, at least over some duration of the infection (Figure 2d). This would give rise to the expectation of considerable levels of viral reassortment within infected hosts, consistent with findings from a guinea pig study that found robust reassortment *in vivo* between phenotypically neutral strains that differed from one another only by silent mutations [18]. In contrast, however, analysis of viral sequence data from a human challenge study indicated very limited effective reassortment, perhaps because of multiple initiating foci of infection [34].

In addition to its effects on viral population dynamics, viral overdispersion across target cells would have important evolutionary consequences. First, reassortment between genetically and phenotypically distinct strains could bring together beneficial mutations on different gene segments or allow for a more effective purging of deleterious mutations. Second, viral overdispersion effectively produces ‘collective infectious units’ [29]. The existence of these collective infectious units will put selection pressures on a virus to evolve cooperative traits, or, conversely, non-cooperative traits that would allow a virus to ‘cheat’. In either case, the importance of quantifying cellular multiplicity of infection is clear, as multiplicity of infection will determine the distribution of viral group sizes, which would in turn affect the types of ‘social interactions’ experienced by viral populations [30]. The within-host macroparasite models presented here provide an approach for estimating the degree of viral overdispersion from fits to viral data. More generally, these models allow for the reality of cellular coinfection dynamics to be integrated into within-host disease models, in a scalable, low-dimensional fashion. While their general formulation has been developed here, these models require assumptions to be made between cellular input and various cellular phenotypes. Empirical studies examining the structure of these relationships is the next critical step to the continued development of these models, and towards their use in better understanding the within-host and evolutionary dynamics of viral infections.

